# Demographic consequences of climate variation along an elevational gradient for a montane terrestrial salamander

**DOI:** 10.1101/130922

**Authors:** Nicholas M. Caruso, Leslie J. Rissler

**Author notes:** NMC ORCID ID# 000-0002-1059-6907.

## Abstract

Climate change represents a significant threat to amphibians, which are already imperiled. However, for many species, the relationship between demographic vital rates (survival and growth) and climate is unknown, which limits predictive models. Here we describe the life history variation of *Plethodon montanus* using capture-recapture data over a period of four years, at five sites along an elevational gradient to determine how survival and growth vary with temperature, precipitation, and how these relationships vary with elevation. We used a hierarchical model to estimate asymptotic size and growth rate, and used a spatial Cormack-Jolly-Seber model to estimate probability of capture and survival, as well as dispersal variance. Our results show that during the active season, growth and survival rates are both positively affected by precipitation, while survival was positively affected by temperature at all elevations, the relationship between growth rates and temperature varied along the elevational gradient. Generally at lower elevations, higher temperatures led to a decrease in growth while at higher elevations the opposite was true. During the inactive season we found elevational variation in the relationship between survival and the amount of snow; at low elevations snowfall was low but survival decreased with increasing snowfall while at higher elevations increasing snowfall lead to higher survival. Our results demonstrate that understanding how the environment can affect salamander demography to develop mechanistic models, will require knowledge of the actual environmental conditions experienced by a given population as well as an understanding of the overall differences in climate at a given site.

## Introduction

Amphibians are one of the most endangered vertebrate taxa (McCallum 2007; Hoffman et al. 2010; IUCN 2016) and face multiple onslaughts including emerging infectious diseases, habitat loss, invasive species, and climate change (Hoffman et al. 2010; Blaustein et al. 2011; Grant et al. 2016). Currently, at least 41% of the approximately 6,500 recognized amphibian species are considered threatened (Hoffman et al. 2010; IUCN 2016) and at least 50% of all salamander species are currently listed as “critically endangered”, “endangered”, or “vulnerable” (IUCN 2016). Trends in declining salamander populations have recently become both taxonomically and geographically widespread (e.g., Rovito et al. 2009; Adams et al. 2013; Spitzen-van der Sluijs et al. 2013). These declines are especially concerning because salamanders represent a significant portion of the total forest biomass and function as keystone predators (Burton and Likens 1975; Milanovich and Peterman 2016).

Given that many populations are already experiencing declines, future changes in climate, represent a compounding threat to amphibian populations (Milanovich et al. 2010; Sutton et al. 2015; Caruso et al. 2017). Recent evidence suggests that contemporary changes in climate have already affected amphibian life history traits (e.g., Reading 2007; Caruso et al. 2014; but see Connette et al. 2015). In addition, warmer temperatures result in metabolic depression (Catenazzi 2016) and slower growth rates of salamanders (Muñoz et al. 2016), which can negatively affect fitness. Under future climate change, populations may become further isolated to higher, cooler elevations (Bernardo and Spotila 2006; Bernardo et al. 2007; Gifford and Kozak 2012; Lyons et al. 2016). Current model predictions of how changes in climate may affect salamander distributions are generally limited to correlative models (e.g., Milanovich et al. 2010; Sutton et al. 2015; Caruso et al. 2017), which do not take into account metrics of demographic vital rates (i.e., survival, growth, and reproduction) as they are lacking for many species. Therefore, current models likely underestimate the effects of future changes in climate (Buckley et al. 2010, Urban et al. 2016).

Demographic vital rates can vary across spatial gradients, and these rates are driven by the biotic (e.g., competition) and the abiotic (e.g., temperature) environment. Lower quality environmental conditions can limit a species’ distribution, while higher quality environments allow for persistence (Gaston 2003). In general, pole-ward range limits are thought to be set primarily by abiotic factors and equator-ward limits by biotic interactions (Schemske et al. 2009). Contrastingly, correlative niche models suggest that amphibian ranges may be more limited at the warmer range edges by the abiotic environment (Cunningham et al. 2016). Although data for montane salamander species are sparse, physiological constraints (Bernardo and Spotila 2006; Gifford and Kozak 2012; Lyons et al. 2016) and results of reciprocal transplant experiments (Cunningham et al. 2009; Caruso et al. 2017) support this trend. However, detailed sampling of vital rates across multiple populations distributed across a species’ range is time-consuming and labor-intensive; therefore, few studies on salamanders have used this information to inform models of range limits and shifts.

As global climates continue to shift, demographic vital rates have become increasingly important to characterize the health of natural populations and to develop informed population models (Pauly 1995; Caswell 2000; Sarrazin and Legendre 2000; Tenhumberg et al. 2004; Coulson et al. 2005; Buckley et al. 2010; Urban et al. 2016). Unfortunately, vital rates and life history traits are unknown for many plethodontid species, adding further uncertainty to their potentially bleak future (e.g., Milanovich et al. 2010). Even when such studies are done, sampling biases such as unobservable ecological states, imperfect and variable detection, and measurement error can distort vital rate estimates of natural populations (Leberg et al. 1989; Royle and Dorazio 2008; Schwarz and Runge 2009; Eaton and Link 2011; Kéry and Schaub 2012; Connette and Semlitsch 2015; Connette et al. 2015; Kéry and Royle 2016). Capture-recapture (CR) methods offer a solution for accounting for these biases; individual observable ecological states can be tracked, while uncertainty in these states can be modeled (Kéry and Schaub 2012; Kéry and Royle 2016). Survival is often a focus of CR studies, as understanding survival, its variation (both temporal and spatial), and the abiotic and biotic factors that drive this variation, are necessary to understanding the underlying spatial and temporal variation in population growth (Lebreton et al. 1992; Saether and Bakke 2000). Similarly, growth is useful for understanding population demographics since larger body size in many species, especially amphibians, is associated with higher fitness (e.g., Petranka 1998). Both survival and growth estimates can be improved using CR methods. Survival can be modeled by accounting for capture probability and dispersal (Lebreton et al. 1992; Schaub et al. 2004; Schaub and Royle 2014), while estimating measurement error and the variation in growth within and among individuals can improve growth estimates (Eaton and Link 2011; Link and Hesed 2015).

In this study, we collected four years of capture-recapture data for *Plethodon montanus* at five sites along an elevational gradient and used hierarchical models to explore the relationship between demography and climate. The objectives of our study were to 1) determine how demographic vital rates (growth and survival) vary along an elevational gradient and among seasons, 2) determine how environmental conditions (temperature and precipitation) affect variation in growth and survival, and 3) determine how the relationship between both survival and growth and environmental conditions varies along the elevational gradient and among seasons. We hypothesized that *P. montanus* vital rates would be driven by climate, whereby warmer and drier conditions would decrease both survival and growth and that lower elevation populations, by virtue of being warmer and drier, will show reduced survival and growth compared to those at higher elevations.

## Methods

### Locations and Sampling

We established five sites along an elevational gradient within the range of *P. montanus* in Pisgah National Forest in 2013: SPG (Spivey Gap; 996m), IMG (Iron Mountain Gap; 1,134m), HG, (Hughes Gap; 1,231m), BBT (Big Butt Trail; 1,300m), and CG (Carver’s Gap; 1,464m). These sites were chosen to minimize the differences among sites in leaf litter depth, aspect, canopy coverage, and number of surface retreats while establishing an elevational gradient within distribution of *P. montanus*.

Within each site, we delineated one-150 m^2^ plot (10 x 15 m). Starting in 2014, we established a grid (25-2 x 3 m sections) within the plot to determine the location of each individual salamander within 0.5 m. Although surveys differed in number of people, effort, and type (i.e., diurnal and nocturnal), salamanders were processed similarly regardless of survey type, amount of effort, or number of surveyors. We captured all salamanders encountered and measured their body size from the snout to the posterior margin of the vent (SVL). We marked *P. montanus* using Visual Implant Elastomer (VIE; Northwest Technology Inc., Shaw Island, Washington) tags, which have minimal effects on fitness and low incidence of tag loss (Bailey 2004). After all salamanders were processed on a particular sampling occasion, we released all individuals back to the original point of capture. See appendix A for additional detail about sampling and site characteristics.

### Site- and Survey-Specific Climate

First, we defined two seasons based on our sampling; the active season (i.e., when salamanders were typically active and when we conducted surveys – 27 May to 13 October), and the inactive season (14 Oct to 26 May). Next, for the length of our study, we obtained daily temperature and precipitation data from the DAYMET database (http://www.daymet.org; Thornton et al. 1997) as covariates in our growth and survival models. We defined temperature for both the active and inactive season as the mean maximum temperature (°C) during the sampling interval, while we defined precipitation for the active season as mean precipitation (mm) and for the inactive season as the mean snow water equivalent (mm) during the sampling interval.

### Asymptotic Size and Growth

To test the degree to which growth is driven by climate, we used a hierarchical model to estimate the effects of season, elevation, temperature, and precipitation on growth rates. Our model estimates the expected size of the *i*th individual at the *t*th time (*ES*_*it*_) using the von Bertalanffy growth curve, parameterized for unknown ages (Fabens 1965) as a function of its expected size at the previous measurement time (*TS*_*it*-1_), elevation-specific asymptotic size (*a*_*elev*_), active [*k(A*)_*it*_] and inactive [*k(IA*)_*it*_] season growth rates scaled for 365 day increments, and the interval between captures (number of days) during the active [Δ*t(A*)_*it*_] or inactive [Δ*t(IA*)_*it*_] seasons (Equation 1).

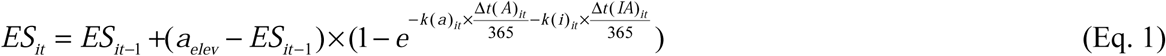

The logarithm of active and inactive season growth rates were subsequently defined by season-specific intercepts [*α*_*k*(*A*)_,*α*_*k*(*IA*)_], as well as covariates for elevation [*βelev*_*k*(*A*)_, *βelev*_*k*(*IA*)_], temperature [*βtemp*_*k*(*A*)_, *βtemp*_*k*(*IA*)_], precipitation [*βprecip*_*k*(*A*)_], SWE [*βswe*_*k*(*IA*)_] and the interaction between elevation and temperature [*βet*_*k*(*A*)_, *βet*_*k*(*IA*)_], precipitation [*βep*_*k*(*A*)_], or SWE [*βes*_*k*(*IA*)_; Equation 2].

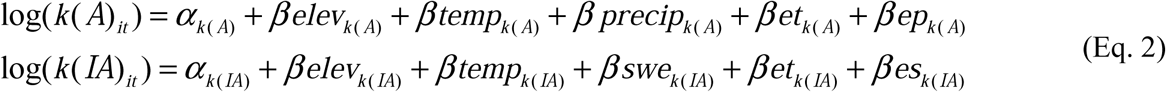

Lastly, our measurements of a given individual (*SVL*_*i,t*_) are described by independent normal random variables with a mean of the expected size (*ES*_*i,t*_) and an estimated standard deviation (σ_SVL_; i.e., measurement error). Therefore, using this hierarchical model we estimated 18 parameters. For further model details and code, see appendix B.

### Bayesian Growth Analysis

To evaluate our growth models, we assigned vague normal priors (mean = 0; variance = 100) to all growth rate covariates, uniform priors for elevation specific asymptotic size (min = 1 50, max = 80), and a vague Gamma prior (shape and rate = 0.001) to the parameter 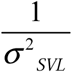 (i.e., precision). All continuous covariates were first scaled and centered prior to model fitting. We fit the model using Markov chain Monte Carlo (MCMC), generating three chains, each with 400,000 iterations. We used an adaptation phase of 1,000, discarded 250,000 burn-in iterations, and used a thinning rate of 50, retaining 3,000 iterations from each chain to estimate posterior distributions (9,000 total samples).

### Capture, Dispersal, and Survival

To test the degree to which survival is driven by climate, we used a spatial Cormack-Jolly-Seber (s-CJS) model (Schaub and Royle 2014) to estimate the effects of season, elevation, temperature, and precipitation on survival while accounting for variation in capture probability, and dispersal. For each individual, we modeled survival to each primary period after its initial capture. Therefore, an individual’s ecological state during the primary period where it is first captured and marked is known (i.e., equal to one). For subsequent primary periods, the ecological state of the *i*th individual at the *t*th primary period (*z*_*i,t*_) is described by a Bernoulli distribution where the probability of success (i.e., the individual is alive, given that it was alive previously) is the product of the probability of survival of the *i*th individual to the *t*th primary period (*ϕ*_*i,t*_) and the ecological state of the *i*th individual at the previous primary period (*z*_*i,t-1*_). Our observation process is likewise described by a Bernoulli distribution where the probability of success (i.e., finding the *i*th, individual, at the *t*th primary period, and *tt*th secondary period, given that it is alive and within the bounds of the study area) is the product of the capture probability (*p*_*i,tt,t*_), ecological state (*z*_*i,t*_), and spatial state (*r*_*i,tt,t*_) of the *i*th individual, at the *t*th primary period, and *tt*th secondary period (Schaub and Royle 2014; Kéry and Schaub 2012).

To account for the fact that some individuals emigrated and thus represent apparent survival (Schaub and Royle 2014), we included each individual’s spatial location within each study site and estimated dispersal from subsequent recaptures. The spatial state (*r*_*i,tt,t*_) of the *i*th individual, at each *tt*th secondary period, and *t*th primary period, is given a value of one if the location in the x- and y-axes (*Gx*_*i,tt,t*_, *Gy*_*i,tt,t*_ respectively) of that individual, at that time is within the study area, while the spatial state receives a value of zero if that individual, at that time is outside the study area. Because we sampled secondary periods within primary periods (2013-2014), we first describe the primary period center of activity in the x- and y-axes (*Gx*_*i,t*_, *Gy*_*i,t*_ respectively). The initial primary period center of activity of the *i*th individual, at the *t*th primary period is described by a uniform distribution, which is bounded by the lower and upper bounds of the x- and y-axes of the plot area (i.e., an individual must be within the study area to be marked). For subsequent primary periods (*t*+*1*), the center of activity of the *i*th individual is normally distributed where the mean is the center of activity at the previous primary period in the x- and y-axes and two estimated precision parameters 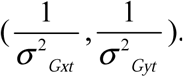. Lastly, the spatial location of the *i*th individual at the *tt*th secondary period and *t*th primary period are also normally distributed where the mean is the center of activity during the *t*th primary period and two estimated precision parameters 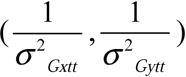. We therefore estimated four parameters for the dispersal portion of our model.

We modeled the logit of the capture probability (*p*_*i,tt,t*_) for the *i*th individual, at the *t*th primary period, and *tt*th secondary period as a function of an intercept (*α*_*p*_), survey type (diurnal or nocturnal; *βsurv*_*p*_), effort (1 or 2; *βeff*_*p*_), number of people (1 or 2; (*β pers*_*p*_), linear and quadratic terms for Julian day (*βjday*_*p*_, 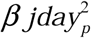 respectively), site (*βsite*_*p*_) and random intercepts for individuals (*ɛ*_*i*_) and primary period (*γ*_*i,t*_; Equation 3).

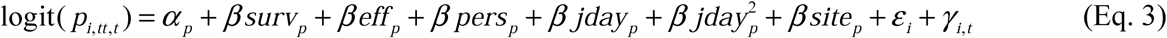

Therefore, for capture probability we estimated twelve total parameters (i.e., ten for the fixed effects and one for the precision component for each of the two random intercepts). Lastly, we modeled the logit of survival (*ϕ*_*i,t*_) of the *i*th individual at the *t*th primary period as a function of an intercept (*α*_*ϕ*_), size (last SVL measurement; *βsize*_*ϕ*_), elevation (*βelev*_*ϕ*_), season (*βseason*_*ϕ*_), temperature, precipitation, and SWE during the active [*βtemp*_*ϕ*(*A*)_, *βprecip*_*ϕ*(*IA*)_] and inactive [*βtemp*_*ϕ*(*A*)_, *βswe*_*ϕ*(*IA*)_] seasons, as well as interactions between elevation with season (*βese*_*ϕ*_), active season temperature and precipitation [*βet*_*ϕ*(*A*)_, *βet*_*ϕ*(*IA*)_] and inactive temperature and precipitation and SWE [*βep*_*ϕ*(*A*)_, *βesw*_*ϕ*(*IA*)_; Equation 4].

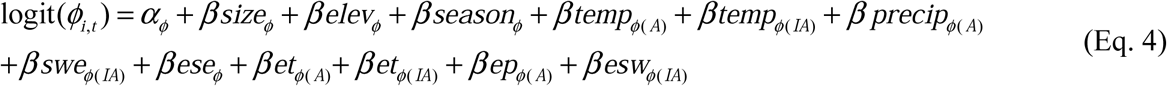

We therefore estimated a total of 13 parameters for survival. For further model details and code, see appendix B.

### Bayesian Survival Analysis

To evaluate our survival models, we first scaled and centered all continuous fixed. We assigned uniform priors (min = 0, max = 10) to all spatial variance estimates. For fixed parameters, we assumed vague normal priors (mean = 0; variance = 100), and random intercepts (*γ*_*t*_ and *ε*_*i*_) were given normal priors, which had estimated precision parameters from a uniform distribution (min = 0; max = 10). We fit the model using MCMC, generating three chains, each with 600,000 iterations. We used an adaptation phase of 1,000, discarded 300,000 burn-in iterations, and used a thinning rate of 50, retaining 6,000 iterations from each chain to estimate posterior distributions (18,000 total samples).

All analyses were performed in program R version 3.3.1 (R Core Team 2016). We used the *jagsUI* package (Kellner 2017) to call JAGS (Plummer 2003), from Program R for MCMC analyses. We examined traceplots of parameters for adequate mixing among chains and the 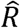 statistic (Gelman et al. 2004) to evaluate model convergence (see appendix C), and we evaluated parameter significance based on the overlap of 95% highest posterior density (HPD) with zero.

## Results

Over 190 diurnal (n = 58; 31%) and nocturnal (n = 132; 69%) surveys, we captured 2,962 salamanders representing nine species (*P. montanus* = 2,413, 81%; non-target species = 549, 19%). We marked a total of 1,343 individuals, and recapture events constituted 1,070 (44%) of our total captures of *P. montanus* captures; we recaptured 559 (42%) individuals at least once (range = 1 – 15 times). Recapture rates generally increased throughout the duration of this study and average recapture rates were at least 40% during the final year of this study at all sites (Appendix A7).

### Growth

We used animals that were captured at least twice for all growth analyses, which included 1,586 total measurements (544 unique individuals), with a range of 92 – 728 measurements per site (36 – 215 unique individuals per site). Although the highest elevation site (CG) had the largest asymptotic size estimate (62.2 mm) this size was similar to the asymptotic size estimate for IMG (62.1 mm), which is approximately 300 meters lower in elevation, while the remaining three sites had smaller asymptotic sizes (Fig. 1A). During the active season, we found significant effects of average precipitation (*βprecip*_*k*(*A*)_; 95% HPD = 0.15 – 0.45) and the interaction between average temperature and elevation (*βet*_*k*(*A*)_; 95% HPD = 0.10 – 0.32; Fig 1B). At all sites, increased precipitation was associated with higher growth rates at all sites, whereas the relationship between temperature and growth rate varied along the elevational gradient; higher temperatures at lower elevations resulted in lower growth rates while the opposite was observed at higher elevations (Fig 2A). Average growth rates during the inactive season were considerably lower (*α*_*k*(*IA*)_; 95% HPD = −28.37 - −6.26) than those during the active season (*α*_*k*(*A*)_; 95% HPD = 0.11 - 0.37) and inactive season growth rates were not significantly influenced by climate (Fig 1C).

**Fig. 1.**
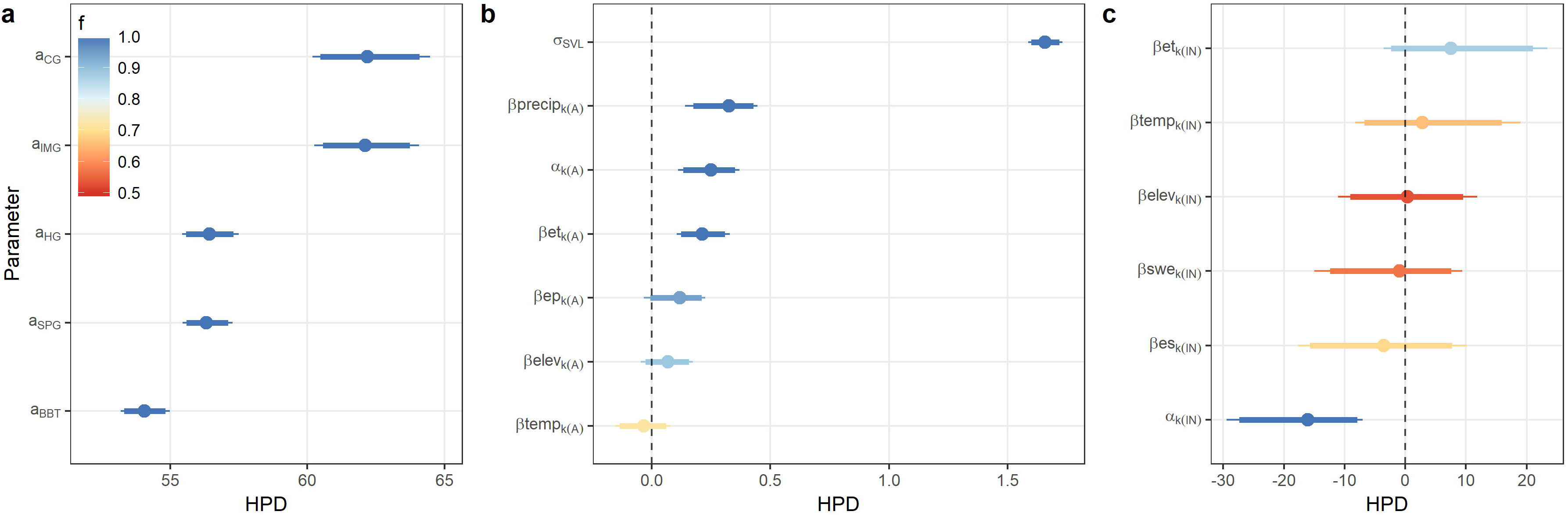
Highest posterior density for growth model with asymptotic size (**a**), active season parameters and measurement error (**b**), and inactive season parameters (**c**). Points represent mean estimates; thick lines show 90% HPD while thin lines show 95% HPD. Colors denote the proportion of the posterior sample that has the same sign as the mean estimate (f).

**Fig. 2.**
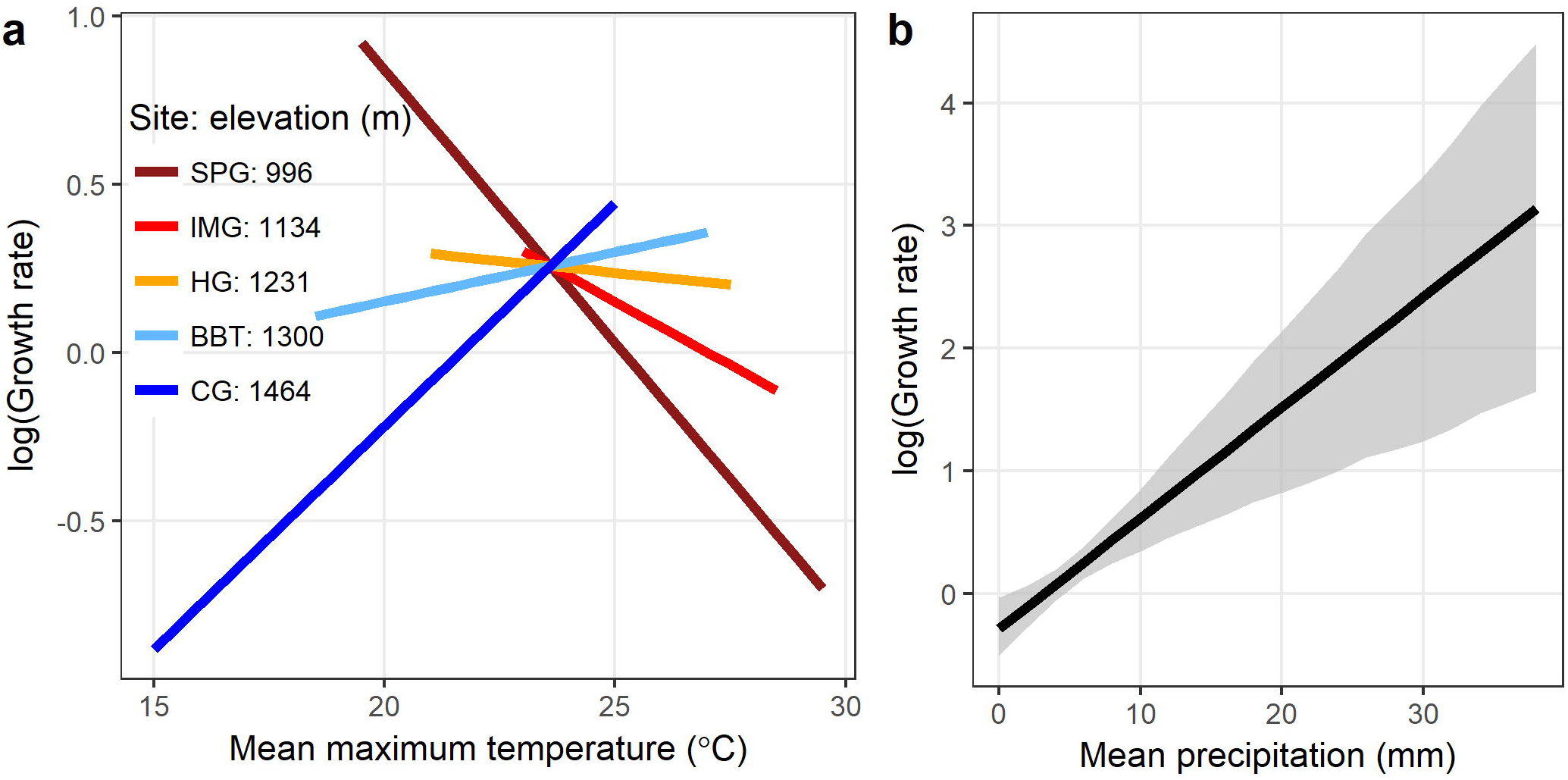
Relationship between the logarithm of growth rate with temperature and elevation (**a**), and precipitation (**b**). Line colors in show site elevation (**a**) while the gray shaded ribbon shows the 95% HPD (**b**). Predicted relationships were limited to the actual range of climate experience by each site (**a**) or at all sites (**b**).

### Dispersal and Capture

Dispersal variance estimates from s-CJS models were similar for primary and secondary seasons with values of approximately one in both the x- and y-axes (Fig. 3A). Therefore, we would expect that 95% of an individual’s movements in the x- and y-axes would be found within ~2 meters from their previous point of capture. Individual variation in capture probability (σ_*ε*_) was lower (95% HPD = 0.01 - 0.20) compared to primary period variation (σ_*γ*_; HPD 95% = 0.67 - 1.04) and we found that survey type (*βsurv*_*p*_; 95% HPD = 2.01 - 2.42), the amount of effort (*βeff*_*p*_; 95% HPD = 0.81 - 1.95), the number of people (*β pers*_*p*_; 95% HPD = 0.41 - 1.18) significantly explained capture probability (Fig. 3B). Capture probability was higher during nocturnal surveys and those that had two people and an increased effort (Fig. 3B).

**Fig. 3.**
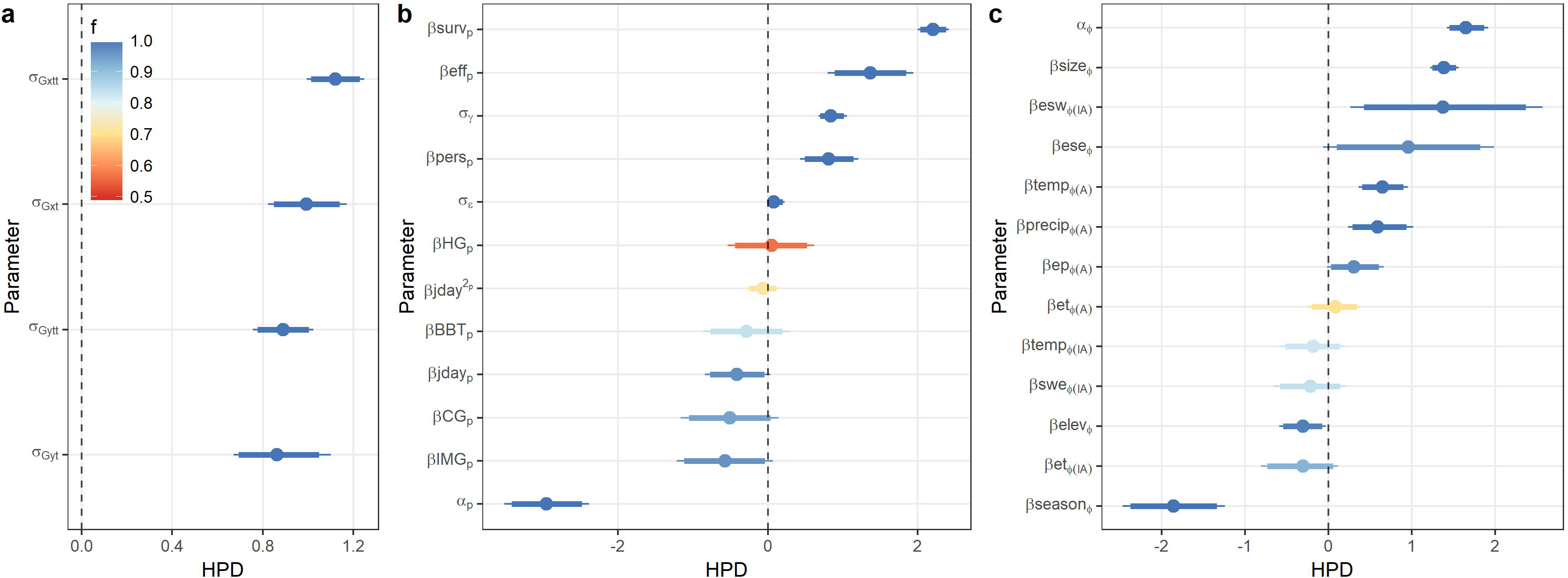
Highest posterior density for s-CJS model with dispersal parameters (**a**), capture parameters (**b**), and survival parameters (**c**). Points represent mean estimates; thick lines show 90% HPD while thin lines show 95% HPD. Colors denote the proportion of the posterior sample that has the same sign as the mean estimate (f).

### Survival

We found that survival was significantly affect by size (*βsize*_*ϕ*_; 95% HPD = 1.22 - 1.56), elevation (*βelev*_*ϕ*_; 95% HPD = −0.58 - −0.02), and season (*βseason*_*ϕ*_; 95% HPD = −2.48 - −1.25; Fig. 3C). Survival increased with increased SVL, was lower at high elevations compared to low elevations, and was higher during the active season compared to the inactive season (Figs. 3C, 4). During the active season both temperature (*βtemp*_*ϕ*(*A*)_; 95% HPD = 0.36 – 0.94) and precipitation (*βprecip*_*ϕ*(*A*)_; 95% HPD = 0.21 – 0.97) positively influenced survival, increased precipitation and higher temperatures were associated with higher survival (Fig. 4A, B). However, during the inactive season we found that survival was significantly influenced by the interaction between average SWE and elevation (*βesw*_*ϕ*(*IA*)_; 95% HPD = 0.22 – 2.51; Fig. 3C). At higher elevations, a greater amount of SWE was associated with higher survival while the opposite was observed at lower elevations (Fig. 4C).

**Fig. 4.**
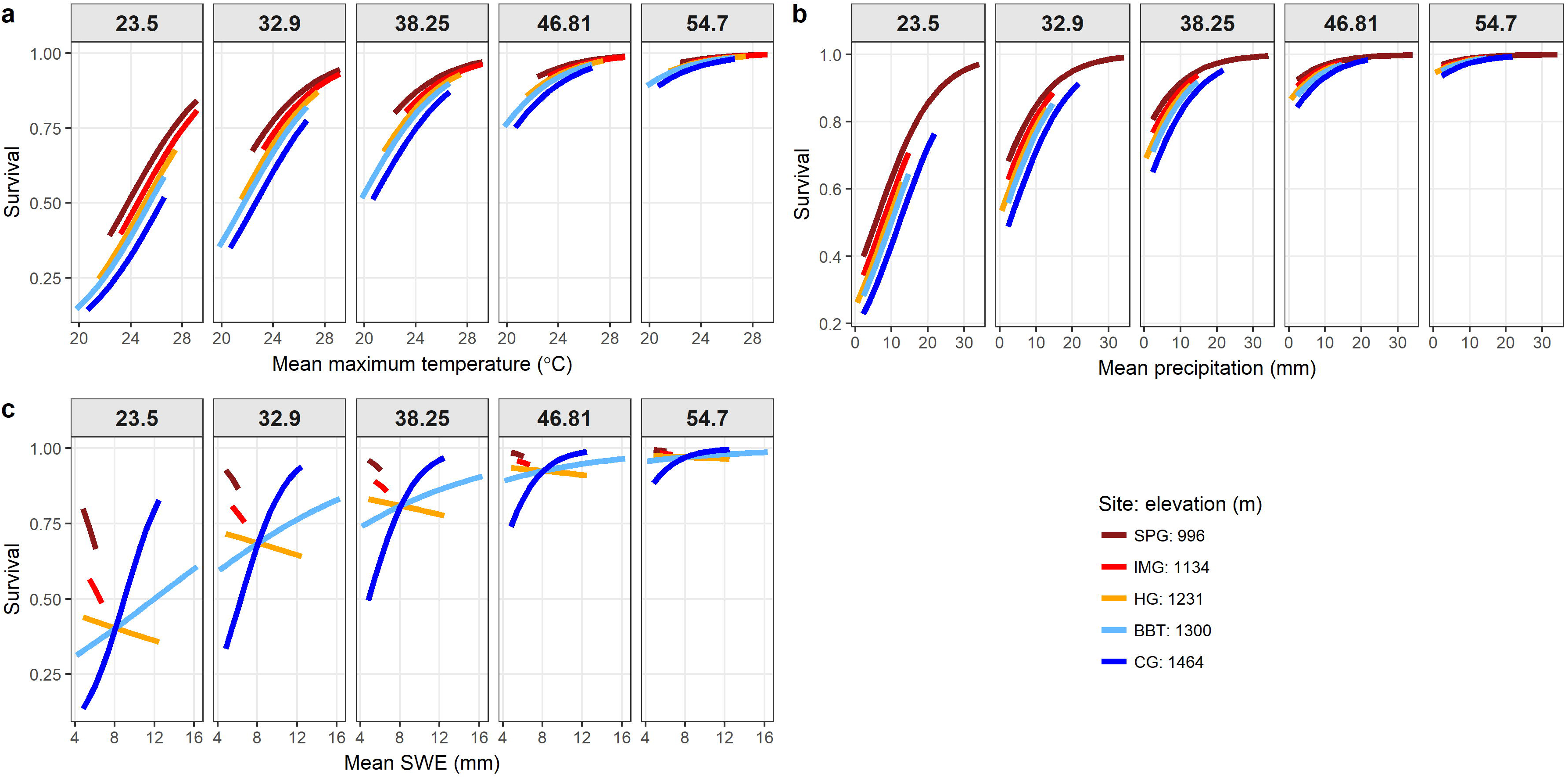
Relationship between survival with temperature (**a**), and precipitation (**b**) and snow water equivalent (SWE; **c**). In all plots, line colors in show site elevation, while the plot facets represent the 10, 30, 50, 70, 90% quantiles of SVL (mm) to show the variation in survival over the range of sizes. Predicted relationships were limited to the actual range of climate experience by each site (**a, b, c**).

## Discussion

We present four years of capture-recapture data to provide a detailed account of how survival and growth of *Plethodon montanus* varies along an elevational gradient and with climate. We found that survival and growth of *P. montanus* is influenced by climate and importantly that the relationship between either survival or growth and climate can vary along an elevational gradient. Our results suggest that *P. montanus* exhibits variation in life history along this elevational gradient, likely resulting from the differences in abiotic environment experienced by those populations. Therefore, understanding how the environment can affect salamander populations, via survival and growth of individuals, may require more than just knowledge of the actual environmental conditions experienced by a given population but also understanding the overall differences in climate at a given site.

### Dispersal

Previous studies have found small home ranges and low dispersal in other species of terrestrial plethodontids (Kleeberger and Werner 1982; Mathis 1991; Merchant 1972; Muñoz et al. 2016). Our observations for *P. montanus* are consistent with these patterns. Dispersal and subsequent immigration can buffer sink populations from declines even when climate negatively affects demographic vital rates and population growth (Brown and Kodric-Brown 1977; Dias 1996; Pulliam 1988; Tavecchia et al. 2016). Unfortunately, montane salamanders, like *P. montanus*, are also physiologically restricted at lower elevations, and tracking suitable climate would likely be limited through warmer valleys and across latitudes, which may exacerbate population isolation and range contractions (Bernardo and Spotila 2006; Kozak and Wiens 2006). The low dispersal observed for *P. montanus* and other terrestrial plethodontids (Cabe et al. 2007; Liebgold et al. 2011; Marsh et al. 2004; Ousterhout and Liebgold 2010; Peterman and Semlitsch 2013) likely further increases their risk of population decline under future climate change.

### Active Season

We found that increased precipitation is associated with both higher growth rates across the elevational gradient (Fig 2B). It has been well-established that rainfall can influence surface activity of plethodontid salamanders (Petranka and Murray 2001; Bendik and Gluesenkamp 2013; Connette et al. 2015), which require moist conditions due to their high rates of evaporative water loss (Spight 1968; Spotila 1972; Spotila and Berman 1976). Conditions, such as increased precipitation, allow for a greater surface activity window and subsequently increased foraging time (Feder and Londos 1984) or may increase the encounter rates of salamanders with their prey (Jaeger 1978, 1980); both of which would likely increase the total number of prey captured and would lead to increases in growth rates, assuming other environmental conditions are equal (e.g., temperature; Catenazzi 2016). Higher precipitation was also associated with an increase in survival across the elevational gradient (Fig 4B). Direct mortality from reduced rainfall is unlikely as salamanders are able to find and take advantage of moist microhabitats in seemingly unfavorable habitats (e.g., Yanev and Wake 1981); however, dehydration can cause a reduction in locomotor performance (Feder and Londos 1984), which may in turn increase predation and decrease survival.

Temperature-sensitivity in various aspects of growth rates (e.g., rates of assimilation and digestion) has been previously demonstrated in plethodontids (e.g., Fitzpatrick 1973; Bobka et al. 1981; Muñoz et al. 2016; Clay and Gifford 2017). In our study, the effect of temperature on growth rates of *P. montanus* varied along the elevational gradient; higher temperatures resulted in lower growth rates at low elevations and higher growth rates at high elevations (Fig. 2A). This pattern may be explained, at least in part, by the range of average temperatures experienced by these populations; high elevation populations generally experienced temperatures lower than those experienced by low elevations (Appendix A4; Fig. 2A). Indeed, Clay and Gifford (2017) found a similar pattern, in which energy assimilation under laboratory conditions in *P. montanus* steadily increased with increasing temperatures but rapidly dropped after reaching an optimum (22.8°C for high elevations and 22.6°C for low elevations) similar to the average temperature at which the growth rate temperature relationship in our study (~23.5 °C) changed from positive (lower temperatures) to negative (higher temperatures; Fig. 2A). Survival, on the other hand, was affected by temperature similarly across our elevational gradient; higher temperatures were associated with higher rates of survival (Fig. 4A). While high temperatures can invariably lead to death in plethodontids, temperatures experienced by our populations never reached higher than 30°C, which is lower than the critical thermal maximum of similar species (Spotila 1972). Increased temperature can have a positive influence on terrestrial salamander endurance and speed (Else and Bennett 1987; Johnson et al. 2010), which may allow them to more readily escape predation. Moreover, under extended warmer conditions, plethodontids may reduce surface activity (Spotila 1972), which would result in a reduction in encounters with potential surface predators. If higher temperatures increased survival via a reduction in surface activity, we would also expect populations to show a reduction in growth rates due to reduced foraging opportunities, which is supported in our growth analyses (Fig. 2A).

### Inactive Season

Our results suggest a disparity in both growth and survival along elevations during the inactive season compared to the active season (Figs. 1 and 3). Low growth rates during the inactive season is likely due to the extended periods of inactivity and lower prey availability, although salamanders have been found on the surface during the winter when temperatures are above freezing (e.g., Catenazzi 2016). Reduced food intake due to the decreased foraging time and/or lower prey availability may also explain the lower survival that we observed during the inactive season compared to the active season. Salamanders that are active during warmer winter conditions may not be able to find the necessary food sources to compensate for this increased activity, leading to decreased body condition and lower survival (Reading 2007; Sinclair et al. 2013; Catenazzi 2016).

During the inactive season, we found that the interaction between SWE and elevation was a significant predictor of survival for *P. montanus* (Fig. 4C). Increased SWE lead to an increase in survival at higher elevations but a decrease in survival at lower elevations; although SWE at the lowest elevations was low and was less variable (Fig. 4C). We suggest two mechanisms by which increases in the amount of snow at higher elevations could increase survival. First, salamanders may have increased survival in areas with more snowpack throughout the winter because of their reduced surface activity, which leads to a reduction in the number of encounters with surface predators (Turbill et al. 2011). Alternatively survival may increase with an increase in snowfall because snow acts as a soil insulator (Decker et al. 2003) and less snowpack can lead to more variable and colder soil temperatures (Groffman et al. 2001; Henry 2008; Bale and Hayward 2010; Brown and DeGaetano 2011). Therefore, hibernating salamanders in areas with more snowpack would have a greater buffer from subzero temperatures (Decker et al. 2003).

Importantly, predictions of salamander population growth under future climate change that only account for responses to the active season conditions may underestimate losses. Warming is predicted to be unequal among seasons, winter months will likely see a greater increase in temperatures than the other seasons (Xia et al. 2014). For logistic reasons, studies of terrestrial plethodontids have typically focused on the active season (i.e., when individuals are available for capture). Therefore, determining the effect of winter conditions on salamander demography (e.g., survival) through the experimental manipulation of temperature or snowpack would improve mechanistic predictive population models (Sanders-DeMott and Templer 2017).

### Conclusions

Future warming is predicted to be a major challenge for Appalachian salamanders (Milanovich et al. 2010; Sutton et al. 2015; Catenazzi 2016; Caruso et al. 2017). Yet mechanistic population growth models are lacking, due, in part, to the paucity of demographic data for many species. Through surveying multiple populations along an elevational gradient, this study was not only able to estimate survival and growth and their variation with relevant climatic factors, but also we were able to demonstrate that the relationships between salamander demography and climate also vary with elevations. This spatial variation in demographic vital rates, and their relationship with relevant climatic factors, is necessary to model population growth and develop conservation strategies (Caswell 2001; Buckley et al. 2010; McLean et al. 2016; Urban et al. 2016). Furthermore, our study demonstrates the importance of climate variation in life history strategies in *P. montanus* likely resulting from both the overall environmental differences in our elevational gradient as well as the variation in climate experience by these populations over the course of this study.

## Acknowledgements

Thanks to C. Houser, J. Jacobs, and E. Kabay for assistance in the field and to D. Adams, C. Staudhammer, G. Starr, S. Wiesner, S. Kunwor, S. George, and two anonymous reviewers for comments that improved this manuscript. This research was funded through Graduate Research Fellowship, E.O. Wilson Fellowship, and the Herpetologists’ League E.E. Williams Research Grant awarded to NMC.

## References

Adams MJ, Miller DAW, Muths E, Corn PS, Grant EHC, Bailey LL, Fellers GM, Fisher RN, Sadinski WJ, Waddle H, Walls SC (2013) Trends in amphibian occupancy in the United States. PLoS ONE 8:e64347. doi:10.1371/journal.pone.0064347.

Bailey LL (2004) Evaluating elastomer marking and photo identification methods for terrestrial salamanders; marking effects and observer bias. Herpetological Review 35:38–41.

Bale JS, Hayward SAL (2010) Insect overwintering in a changing climate. Journal of Experimental Biology 213:980–994.

Bendik NF, Gluesenkamp AG (2013) Body length shrinkage in an endangered amphibian is associated with drought. Journal of Zoology 290:35–41.

Bernardo J, Spotila JR (2006) Physiological constraints on organismal response to global warming: mechanistic insights from clinally varying populations and implications for assessing endangerment. Biology Letters 2:135–139.

Bernardo J, Ossola RJ, Spotila J, Crandall KA (2007) Interspecific physiological variation as a tool for cross-species assessments of global warming-induced endangerment: validation of an intrinsic determinant of macroecological and phylogeographic structure. Biology Letters 3:695–698.

Blaustein AR, Walls SC, Bancroft BA, Lawler JJ, Searle CL, Gervasi SS (2010) Direct and indirect effects of climate change on amphibian populations. Diversity 2:281–313.

Bobka MS, Jaeger RG, McNaught DC (1981) Temperature dependent assimilation efficiencies of two species of terrestrial salamanders. Copeia 1981:417–421.

Brown PJ, DeGaetano AT (2011) A paradox of cooling winter soil surface temperatures in a warming northeastern United States. Agricultural and Forest Meterology 151:947–956.

Brown JH, Kodric-Brown A (1977) Turnover rates in insular biogeography: effect on immigration on extinction. Ecology 58:445–449.

Buckley LB, Urban MC, Angilletta MJ, Crozier LG, Rissler LJ, Sears MW (2010) Can mechanism inform species’ distribution models? Ecology Letters 13:1041–1054.

Burton TM, Likens GE (1975) Energy flow and nutrient cycling in salamander populations in the Hubbard Brook Experimental Forest, New Hampshire. Ecology 56:1068–1080.

Cabe PR, Page RB, Hanlon TJ, Aldrich ME, Marsh DM (2007) Fine-scale population differentiation and gene flow in a terrestrial salamander (Plethodon cinereus) living in continuous habitat. Heredity 98:53–60.

Caruso NM, Jacobs JF, Rissler LJ (2017). An experimental approach to understanding elevation limits in a montane terrestrial salamander, Plethodon montanus. Submitted to Copeia bioRxiv doi: https://doi.org/10.1101/131573

Caruso NM, Sears MW, Adams DC, Lips KR (2014) Widespread rapid reductions in body size of adult salamanders in response to climate change. Global Change Biology 20:1751–1759.

Caswell H (2000) Prospective and retrospective perturbation analyses: their roles in conservation biology. Ecology 81:619–627.

Caswell H (2001) Matrix Population Models: Construction, Analysis, and Interpretation. 2nd edn. Sinauer Associates, Sunderland

Catenazzi A (2016) Ecological implications of metabolic compensation at low temperatures in salamanders. PeerJ 14:e2072 10.7717/peerj.2072.

Clay TA, Gifford ME (2017) Population level differences in thermal sensitivity of energy assimilation in terrestrial salamanders. Journal of Thermal Biology 64:1–6.

Connette GM, Crawford JA, Peterman WE (2015) Climate change and shrinking salamanders: alternative mechanisms for changes in plethodontid salamander body size. Global Change Biology 21:2834–2843.

Connette GM, Semlitsch RD (2015) A multistate mark-recapture approach to estimating survival of PIT-tagged salamanders following timber harvest. Journal of Applied Ecology 52:1316–1324.

Coulson T, Gaillard J-M, Festa-Bianchet M (2005) Decomposing the variation in population growth into contributions from multiple demographic rates. Journal of Animal Ecology 74:789–801.

Cunningham HR, Rissler LJ, Apodaca JJ (2009) Competition at the range boundary in the slimy salamander: using reciprocal transplants for studies on the role of biotic interactions in spatial distributions. Journal of Animal Ecology 78:52–62.

Cunningham HR, Rissler LJ, Buckley LB, Urban MC (2016) Abiotic and biotic constraints across reptile and amphibian ranges. Ecography 39:1–8.

Decker KL, Wang MD, Waite C, Scherbatskoy T (2003) Snow removal and ambient air temperature effects on forest soil temperatures in northern Vermont. Soil Science Society of America Journal 67:1234–1243.

Dias PC (1996) Sources and sinks in population biology. Trends in Ecology and Evolution 11:326–330.

Eaton MJ, Link WA (2011) Estimating age from recapture data: integrating incremental growth measures with ancillary data to infer age-at-length. Ecological Applications 21:2487–2497.

Else, PL Bennett AF (1987) The thermal-dependence of locomotor performance and muscle contractile function in the salamander Ambystoma tigrinum nebulosum. Journal of Experimental Biology 128:219–233.

Fabens AJ (1965) Properties and fitting of von Bertalanffy growth curve. Growth 29:265–289.

Feder ME, Londos PL (1984) Hydric constraints upon foraging in a terrestrial salamander Desmognathus ochrophaeus (Amphibia: Plethodontidae). Oecologia 64:413–418.

Fitzpatrick LC (1973) Influence of seasonal temperatures on the energy budget and metabolic rates of the northern two-lined salamander Eurycea bislineata bislineata. Comparative Biochemistry and Physiology 45A:807–818.

Gaston KJ (2003) The Structure and Dynamics of Geographic Ranges. Oxford University Press.

Gelman A, Carlin JB, Stern HS, Rubin DB (2004) Bayesian data analysis, 2nd edn. CRC/Capman and Hall, Boca Raton, FL.

Gifford ME, Kozak KH (2012) Islands in the sky or squeezed at the top? Ecological causes of elevational range limits in montane salamanders. Ecography 35:193–203.

Grant EHC, Miller DAW, Schmidt BR, Adams MJ, Amburgey SM, Chambert T, Cruickshank SS, Fisher RN, Green DM, Hossack BR, Johnson PTJ, Joseph MB, Rittenhouse TAG, Ryan ME, Waddle JH, Walls SC, Bailey LL, Fellers GM, Gorman TA, Ray AM, Pilliod DS, Price SJ, Saenz D, Sadinski W, Muths E (2016) Quantitative evidence for the effects of multiple drivers on continental-scale amphibian declines. Scientific Reports 6:25625 doi:10. 1038/srep25625.

Groffman PM, Rustad LE, Templer PH, Campbell JL, Christenson LM, Lany N, Socci AM, Vadebncoeur MA, Schaberg PG, Wilson G, Driscoll C, Fahey TJ, Fisk MC, Goodale CL, Green MB, Hamburg SP, Johnson CE, Mitchell MJ, Morse JL, Pardo LH, Rodenhouse NL (2012) Long-term integrated studies show complex and surprising effects of climate change in the northern hardwood forest. BioScience 62:1056–1066.

Henry HAL (2008) Climate change and soil freezing dynamics: historical trends and projected changes. Climatic Change 87:421–434.

Hoffman M, et al (2010) The impact of conservation on the status of the world’s vertebrates. Science 330:1503–1509.

IUCN (2016) The IUCN red list of threatened species. Version 2016-3. <www.iucnredlist.org>. Downloaded on 22 March 2017.

Jaeger RG (1978) Plant climbing by salamanders: periodic availability of plant-dwelling prey. Copeia 1978:686–691.

Jaeger RG (1980) Fluctuations in prey availability and food limitation for a terrestrial salamander. Oecologia 44:335–341.

Johnson JR, Johnson BB, Shaffer HB (2010) Genotype and temperature affect locomotor performance in a tiger salamander hybrid swarm. Functional Ecology 24:1073–1080.

Kéry M, Royle JA (2016) Applied hierarchical modeling in ecology: analysis of distribution, abundance and species richness in R and BUGS (volume 1 – prelude and static models). Academic Press, San Diego, CA.

Kéry M, Schaub M (2012) Bayesian population analysis using WinBUGS – a hierarchical perspective. Academic Press, San Diego, CA.

Kellner K (2017) jagsUI: a wrapper around ‘rjags’ to streamline ‘JAGS’ analyses. R package version 1.4.9.

Kleeberger SR, Werner JK (1982) Home range and homing behavior of Plethodon cinereus in Northern Michigan. Copeia 1982:409–415.

Kozak KH, Wiens JJ (2006) Does niche conservatism drive speciation? A case study in North American salamanders. Evolution 60:2585–2603.

Leberg PL, Brisbin IL, Smith MH, White GC (1989) Factors affecting the analysis of growth-patterns of large mammals. Journal of Mammalogy 70:275–283.

Lebreton J-D, Burnham KP, Clobert J, Anderson DR (1992) Modeling survival and testing biological hypotheses using marked animals: a unified approach with case studies. Ecological Monographys 62:67–118.

Liebgold EB, Brodie III ED, Cabe PR (2011) Female philopatry and male-biased dispersal in a direct-developing salamander, Plethodon cinereus. Molecular Ecology 20:249–257.

Link WA, Hesed KM (2015) Individual heterogeneity in growth and age at sexual maturity: a gamma process analysis of capture-mark-recapture data. Journal of Agricultural, Biological, and Environmental Statistics 20:343–352.

Lyons MP, Shepard DB, Kozak KH (2016) Determinants of range limits in montane woodland salamanders (genus Plethodon). Copeia 104:101–110.

Marsh DM, Thakur KA, Bulka KC, Clarke LB (2004) Dispersal and colonization through open fields by a terrestrial, woodland salamander. Ecology 85:3396–3405.

Mathis A (1991) Territories of male and female terrestrial salamanders: costs, benefits, and intersexual spatial associations. Oecologia 86:433–440.

McLean N, Lawson CR, Leech DI, van de Pol M (2016) Predicting when climate-driven phenotypic change affects population dynamics. Ecology Letters 19:595–608.

McCallum ML (2007) Amphibian decline or extinction? Current declines dwarf background extinction rate. Journal of Herpetology 41:483–491.

Merchant H (1972) Estimated population size and home range of the salamanders Plethodon jordani and Plethodon glutinosus. Journal of the Washington Academy of Sciences 62:248–257.

Milanovich JR, Peterman WE (2016) Revisiting Burton and Likens (1975): nutrient standing stock and biomass of a terrestrial salamander in the Midwestern United States. Copeia 104:165–171.

Milanovich JR, Peterman WE, Nibbelink NP, Maerz JC (2010) Projected loss of a salamander diversity hotspot as a consequence of projected global climate change. PLoS ONE 5:e12189 DOI 10.1371/journal.pone.0012189.

Muñoz DJ, Hesed KM, Grant EHC, Miller DAW (2016) Evaluating within-population variability in behavior and demography for the adaptive potential of a dispersal-limited species to climate change. Ecology and Evolution 6:8740–8755.

Ousterhout B, Liebgold EB (2010) Dispersal versus site tenacity of adult and juvenile red-backed salamanders (Plethodon cinereus). Herpetologica 63:269–275.

Pauly D (1995) Anecdotes and shifting baseline syndrome of fisheries. Trends in Ecology and Evolution 10:430. doi:10.1016/S0169-5347(00)89171-5

Peterman WE, Semlitsch RD (2013) Fine-scale habitat associations of a terrestrial salamander: the role of environmental gradients and implications for population dynamics. PLoS ONE 8:e62184. doi:10.1371/journal.pone.0062184.

Petranka JW (1998) Salamanders of the United States and Canada. Smithsonian Institution Press, Washington, DC.

Petranka JW, Murray SS (2001) Effectiveness of removal sampling for determining salamander density and biomass: a case study in an Appalachian streamside community. Journal of Herpetology 35:36–44.

Plummer M (2003) JAGS: A program for analysis of Bayesian graphical models using Gibbs sampling. Proceedings of the 3rd International Workshop on Distributed Statistical Computing (DSC2003), pp 20–22.

Pullimam HR (1988) Sources, sinks, and population regulation. The American Naturalist 132:652–661.

R Core Team (2016) R: a language and environment for statistical computing. R Foundation for Statistical Computing, Vienna, Austria.

Reading CJ (2007) Linking global warming to amphibian declines through its effects on female body condition and survivorship. Oecologia 151:125–131.

Rovito SM, Parra-Olea G, Vásquez-Almazán CR, Papenfuss TJ, Wake DB (2009) Dramatic declines in neotropical salamander populations are an important part of the global amphibian crisis. Proceedings of the National Academy of Sciences USA 106:3231–3236.

Royle JA, Dorazio RM (2008) Hierarchical modeling and inference in ecology. Academic Press, San Diego, CA.

Saether B-E, Oyvind B (2000) Avian life history variation and contribution of demographic traits to the population growth rate. Ecology 81:642–653.

Sanders-DeMott R, Templer PH (2017) What about winter? Integrating the missing season into climate change experiments in seasonally snow covered ecosystems. Methods in Ecology and Evolution doi:10.1111/2041-210X.12780

Sarrazin F, Legendre S (2000) Demographic approach to releasing adults versus young in reintroductions. Conservation Biology 14:488–500.

Schaub M, Gimenez O, Schmidt BR, Pradel R (2004) Estimating survival and temporary emigration in the multistate capture-recapture framework. Ecology 85:2107–2113.

Schaub M, Royle JA (2014) Estimating true instead of apparent survival using spatial Cormack-Jolly-Seber models. Methods in Ecology and Evolution 4:1316–1326.

Schemske DW, Mittelbach GG, Cornell HV, Sobel JM, Roy K (2009) Is there a latitudinal gradient in the importance of biotic interactions? Annual Review of Ecology, Evolution, and Systematics 40:245–269.

Schwarz LK, Runge MC (2009) Hierarchical Bayesian analysis to incorporate age uncertainty in growth curve analysis and estimates of age from length: Florida manatee (Trichechus manatus) carcasses. Canadian Journal of Fisheries and Aquatic Sciences 66:1775–1789.

Sinclair BJ, Stinziano JR, Williams CM, MacMillan HA, Marshall KE, Storey KB (2013) Realtime measurement of metabolic rate during freezing and thawing of the wood frog, Rana sylvatica: implications for overwinter energy use. Journal of Experimental Biology 216:292–302.

Spight TM (1968) The water economy of salamanders: evaporative water loss. Physiological Zoology 41:195–203.

Spitzen-van der Sluijs AM, Spikmans F, Bosman W, de Zeeuw M, van der Meij T, Goverse E, Kik M, Pasmans F, Martel A (2013) Rapid enigmatic decline drives the fire salamander (Salamandra salamandra) to the edge of extinction in the Netherlands. Amphibia-Reptilia 34:233–239.

Spotila JR (1972) Role of temperature and water in the ecology of lungless salamanders. Ecological Monographs 42:95–125.

Spotila JR, Berman EN (1976) Determination of skin resistance and the role of the skin in controlling water loss in amphibians and reptiles. Comparative Biochemistry and Physiology 55A:407–411.

Sutton WB, Barrett K, Moody AT, Loftin CS, deMaynadier PG, Nanjappa P (2015) Predicted changes in climatic niche and climate refugia of conservation priority salamander species in the northeastern United States. Forests 6:1–26. doi:10.3390/f6010001.

Tavecchia G, Tenan S, Pradel R, Igual J-M, Genovart M, Oro D (2016) Climate-driven vital rates do not always mean climate-driven population. Global Change Biology 22:3960–3966.

Tenhumberg B, Tyre AJ, Shea K, Possingham HP (2004) Linking wild and captive populations to maximise species persistence: optimal translocation strategies. Conservation Biology 18:1304–1314.

Thornton PE, Running SW, White MA (1997) Generating surfaces of daily meteorological variables over large regions of complex terrain. Journal of Hydrology 190:214–251.

Turbill C, Bieber C, Ruf T (2011) Hibernation is associated with increased survival and the evolution of slow life histories among mammals. Proceedings of the Royal Society B: Biological Sciences 278:3355–3363. doi:10.1098/rspb.2011.0190.

Urban MC, Bocedi G, Hendry AP, Mihoub J-B, Pe’er G, Singer A, Bridle JR, Crozier LG, De Meester L, Godsoe W, Gonzalez A, Hellmann JJ, Holt RD, Huth A, Johst K, Krug CB, Leadley PW, Palmer SCF, Pantel JH, Schmitz A, Zollner PA, Travis JMJ (2016) Improving the forecast for biodiversity under climate change. Science 353:aad8466(2016) DOI: 10.1126/science.aad8466.

Xia J, Chen J, Piao S, Ciais P, Luo Y, Wan S (2014) Terrestrial carbon cycle affected by non-uniform climate warming. Nature Geoscience 7:173–180.

Yanev KP, Wake DB (1981) Genic differentiation in a relict desert salamander, Batrachoseps campi. Herpetologica 37:16–28.

